# Connexin 43 drives glioblastoma cancer stem cell phenotypes through a WNK lysine-deficient protein kinase 1-c-MYC signaling axis

**DOI:** 10.1101/2024.07.01.601549

**Authors:** Erin E. Mulkearns-Hubert, Nicole Hajdari, Ellen S. Hong, Ashley P. Jacobs, Kristen E. Kay, Sabrina Z. Wang, Daniel J. Silver, Christopher G. Hubert, Justin D. Lathia

## Abstract

The coordination of cellular processes such as growth and survival relies on communication between cells through gap junctions. Gap junction intercellular communication is driven by connexin proteins, which also mediate protein-protein interactions and communication with the extracellular space via hemichannels. Despite their essential roles, connexin function in cancer is context dependent, with connexin 43 (Cx43) reported to both promote and suppress tumor growth in glioblastoma, the most common primary malignant brain tumor. Here, we detect expression of Cx43 in glioblastoma patient-derived cancer stem cells and demonstrate that Cx43 is essential for their survival and self-renewal. Mechanistically, depletion of Cx43 reduces c-MYC expression through reduced levels of the upstream mediator WNK lysine-deficient protein kinase 1 (WNK1). Depletion of WNK1 phenocopies Cx43 knockdown and reduces MYC expression and tumor growth. Together, these results define a novel signaling axis downstream of Cx43 that promotes tumor growth and cancer stem cell phenotypes in glioblastoma.

## Introduction

Patients diagnosed with glioblastoma (GBM), the most common primary malignant brain tumor, typically experience a median survival of less than two years after diagnosis^1^. Standard-of-care therapy consisting of maximal safe surgical resection followed by radiation and chemotherapy with the alkylating agent temozolomide has produced a minimal extension in survival, and tumors universally recur. Therapeutic resistance and recurrence of GBM are driven by a combination of tumor-intrinsic characteristics such as the presence of therapeutically resistant, self-renewing cancer stem cells (CSCs), in addition to tumor cell-extrinsic interactions with the tumor microenvironment, including with immune cells. The ability to target these tumor-supportive features will aid in future therapeutic development efforts.

In a normal tissue, adjacent cells communicate directly with one another to coordinate complex processes and also utilize cell-cell communication methods to sense and interact with the surrounding microenvironment (reviewed in ^2,3^). One major method of cell-cell communication, gap junction intercellular communication (GJIC), is mediated by gap junctions composed of connexin proteins. Six connexin proteins form a hexameric structure, termed a connexon or hemichannel, with a central pore that opens to allow small molecules to pass through. Connexons in the membrane of one cell dock with a connexon on an adjacent cell, allowing the transfer of material, such as metabolites, ions, miRNAs, and some proteins smaller than 1 kD, between cells. In some situations, connexons have been shown to function as hemichannels, allowing the movement of small molecules between a cell and the extracellular space^2,4^. Connexins also bind to a number of intracellular proteins, mainly through their C-terminal cytoplasmic tails, and act as scaffolds to link other proteins together. There are 21 connexin proteins in humans, with the main sequence divergence falling within the C-terminal tails. For this reason, specifically targeting connexins has proven challenging.

Connexins and gap junctions have historically been thought of as tumor suppressive, based on reduced GJIC and the frequent loss of connexin expression in tumor tissue, including in GBM^2,5–7^. However, we and others have more recently shown that certain cancer types depend on the expression of specific connexins for both gap junction-dependent and gap junction-independent functions^3^. Specifically, we showed that GBM CSCs from patient-derived xenograft (PDX) models depend on cell-cell communication through connexin 46 (Cx46)^8,9^ and that triple-negative breast CSCs depend on the formation of an intracellular gap junction-independent complex with NANOG and focal adhesion kinase (FAK) by Cx26^10,11^. In both cases, targeting of connexin activity reduced tumor growth and suppressed CSC self-renewal. Although specific targeting of connexins remains a challenge in vivo, the concept of reducing connexin function, either directly or by inhibiting downstream events, may be therapeutically beneficial.

The most widely expressed and frequently studied connexin, Cx43, has been reported to have both tumor promoting and tumor suppressing roles in a number of cancers, including GBM. Expression levels of Cx43 vary widely among GBM patient samples and tumor cell lines and are frequently – but not universally – reported to be lower than those of normal brain^12–16^. Functionally, Cx43 expression in GBM has been shown to drive pro-tumorigenic properties such as therapeutic resistance^17–24^, cell proliferation and survival^25–27^, and tumor cell invasion and migration^27–29^. However, contrasting results have also been reported for Cx43, including reduced proliferation and migration upon introduction of Cx43 into GBM cell lines^30–37^. Many of these studies of Cx43 in GBM have been limited by the use of traditional, serum-cultured cell lines, either human or rat, that do not recapitulate the cellular heterogeneity and molecular alterations observed in patient tumors^38^. Studies in CSC models of GBM, while less abundant, have produced similarly contrasting results; specifically, Cx43 has been shown to suppress tumor growth by inhibiting SRC signaling^33,36,39,40^ but also to promote chemoresistance to temozolomide, the GBM standard-of-care chemotherapy, by activating PI3K/AKT^18,19^.

Here, we identify a role for Cx43 in maintaining GBM CSC proliferation and survival through expression of the c-MYC oncogene. Furthermore, we identify expression of WNK lysine-deficient protein kinase 1 (WNK1) as a signaling intermediate between Cx43 and C-MYC. Together, this work has revealed a novel Cx43-WNK1-MYC signaling axis that drives GBM CSC properties.

## Materials and Methods

### Patient-derived GBM xenografts

The GBM patient derived PDX CSC models T3691, T3832, T387, T4121, and T4302 were obtained from Duke University (originally from Dr. Jeremy Rich) via material transfer agreement^8,41–43^. The DI318 model was generated at Cleveland Clinic, and we have used it previously^44^. BT124 was obtained from the University of Calgary (Dr. Samuel Weiss)^45^. L1 and L2 were obtained from University of Florida (originally from Dr. Brent Reynolds)^46,47^. GBM23 was obtained from MD Anderson Cancer Center (originally from Dr. Erik Sulman) and used previously^48^. The GBM12 model was obtained from the Mayo Clinic (Dr. Jann Sarkaria). All models were de-identified and established in accordance with an Institutional Review Board-approved protocol with informed-consent obtained from patients.

GBM PDX models were cultured in suspension in Petri dishes in Neurobasal medium without phenol red (ThermoFisher Scientific) supplemented with 1% penicillin/streptomycin (ThermoFisher Scientific), 2% B27 without vitamin A (ThermoFisher Scientific), 20 ng/mL EGF (R&D Systems), 20 ng/mL FGF (R&D Systems), 1 mM sodium pyruvate (ThermoFisher Scientific), and 2 mM L-glutamine (Lerner Research Institute Media Core). Cells were subcultured using Accutase (BioLegend) to obtain a single-cell suspension. When adherence was required, cells were incubated with GelTrex (ThermoFisher Scientific) diluted to approximately 1:1000 in the culture media and cultured on tissue culture-treated dishes.

### Additional cell lines

Conventional human cancer cell lines 293T (human embryonic kidney; ATCC: CRL-11268), MDA-MB-231 (breast adenocarcinoma; ATCC: HTB-26), LNCaP (prostate carcinoma; ATCC: CRL-1740), and HeLa (cervical cancer; ATCC: CCL-2) cells were obtained from the Cleveland Clinic Lerner Research Institute Cell and Media Production Core. All cells were maintained in DMEM with 10% FBS and 1% pen-strep in a humidified incubator with 5% CO_2_. Cells were passaged using trypsin. Normal human astrocytes were obtained from Lonza and cultured in supplemented AGM™ Astrocyte Growth Medium (Lonza) for limited passages according to the vendor’s protocols.

### Plasmids and DNA constructs

Predesigned Cx43 and WNK1 shRNA constructs in pLKO.1-puro were purchased from Sigma-Aldrich. Their respective clone ID numbers are as follows: Cx43 sh73 - TRCN0000059773; Cx43 sh76 TRCN0000059776; Cx43 sh77 - TRCN0000059777; WNK1 sh491 - TRCN0000196491; WNK1 sh918 - TRCN0000000918; WNK1 sh920 - TRCN0000000920. The SHC002 Non-Mammalian shRNA Control Plasmid was used as the non-targeting sequence.

### Lentiviral preparation and transduction

Lentivirus was prepared by calcium phosphate transfection of 293T cells using psPAX (Addgene # 12260) and pMD2.G (Addgene # 12259), both produced by Didier Trono. Virus was harvested on days 2 and 3 after transfection, concentrated via polyethylene glycol precipitation, and stored at -80°C until use.

For lentiviral transduction, cells were plated in supplemented neurobasal media as described above with 1:1000 Geltrex (ThermoFisher Scientific) into 6-well plates. Cells were then infected with lentivirus at 5 µL/mL 2 days later. Cells were then selected with 2 µg/mL puromycin (ThermoFisher Scientific) and collected for assays 2-3 days later. Due to the dramatic viability defects observed with shRNA against both Cx43 and WNK1, cells were freshly transduced for each experiment, and any remaining cells were discarded.

### In vivo tumor cell implantation

All animal usage was performed in accordance with protocols approved by the Cleveland Clinic Institutional Animal Care and Use committee. A total of ten 6-week-old mice per group, five male and five female, were implanted with either a non-targeting shRNA, WNK1 sh491, or WNK1 sh920. The mice were anesthetized via 2% isoflurane gas and placed in a stereotaxic instrument. An insulin syringe with a 31-gauge needle was secured to a probe holder, and the needle was passed through the scalp 0.5 mm rostral and 1.8 mm lateral to the bregma. The needle was inserted 3.5 mm underneath the scalp, where 25,000 cells suspended in unsupplemented Neurobasal media were injected slowly into the mice. The needle was held in place for 60 seconds before being slowly removed from the mouse’s scalp. The experiment was blinded to the investigators who injected the mice and to the investigator who measured the endpoints of each mouse.

### Proliferation and apoptosis assays

For each condition, proliferation and apoptosis assays were plated at 2,000 cells per well in triplicate into a 96-well white-walled, clear-bottom plate (BD Bioscience). CellTiter Glo (Promega) was used to measure ATP levels on days 0, 3, and 5 after plating and used as a surrogate for cell viability. Values were normalized to day 0 and then to the non-targeting control to obtain a fold-change value. Apoptosis was measured similarly by determining active caspases 3/7 via CaspaseGlo 3/7 (Promega) on day 3 after plating. Values were normalized to cell number as measured via CellTiter at the same time point.

### Limiting-dilution analysis

GBM CSCs were dissociated using Accutase (BioLegend) and subsequently plated into 96-well suspension plates (Sarstedt) in decreasing cell numbers (20, 10, 5, and 1 cells/well) with 24 replicates for each cell number. After 10-14 days, wells were evaluated for the presence of spheres, which was recorded in a binary manner. The stem cell frequency was subsequently calculated using the online Extreme Limiting Dilution Analysis algorithm (https://bioinf.wehi.edu.au/software/elda/)^49^.

### Immunoblotting and antibodies

To prepare lysates, cells were washed with PBS and lysed with lysis buffer (0.5% NP-40, 10 mM Tris-Cl pH 7.5, 1 mM EDTA pH 8.0, 150 mM NaCl) supplemented 1% phosphatase inhibitors and 1% protease inhibitors (Sigma-Aldrich; P8340 and P5726). Insoluble debris was removed via centrifugation at 21,000 x g for 10 minutes at 4°C. Protein concentration was determined via the Bio-Rad protein assay using a 1 mg/mL solution of bovine serum albumin (BSA) for standardization. For blotting, 15-50 μg of protein lysate was separated via SDS-PAGE electrophoresis. Pre-cast 4-15% gels were used to resolve WNK1 protein. Gels were subsequently transferred onto PVDF membranes (Millipore, IPVH00010), which were blocked for 45 minutes to 1 hour with 5% nonfat milk. Primary antibodies were then incubated on the membranes for 2-7 days at 4°C in 5% BSA in Tris-buffered saline with Tween-20 (TBST) before being washed with TBST. Membranes were then incubated with the appropriate horseradish peroxidase-tagged secondary antibody for 2.5 to 3 hours at room temperature. Primary antibodies used were anti-Cx43 at 1:1000 dilution (Cell Signaling, 3512S), anti-WNK1 at 1:1000 dilution (Cell Signaling, 4979S), anti-c-MYC at 1:1000 dilution (Cell Signaling, 9402S), Na/K ATPase at 1:2000 (ThermoFisher Scientific, MA3-928), and Lamin A/C at 1:3000 (ProteinTech, 10298-1-AP). The membranes were developed using the Pierce ECL 2 Western Blotting Substrate (ThermoFisher Scientific, 80196) and imaged on a Bio-Rad ChemiDoc MP imaging system. Anti-Actin hFAB Rhodamine Antibody (Bio-Rad, 10000068189) was used as loading control.

### Subcellular fractionation

PDX CSC lines DI318, T3691, and T3832 were subjected to subcellular fractionation in accordance with the manufacturer’s instructions for the Subcellular Protein Fractionation Kit for Cultured Cells (ThermoFisher, #78840). Approximately 6 million cells in a packed cell volume of 20 µL was used for all three cell models. Equal volumes of fractionated lysates were subjected to immunoblotting as described above for Cx43, using Na/K ATPase, actin, and lamin a/c as fractionation controls. Membranes were imaged on the Bio-Rad ChemiDoc MP imaging system.

### Enzyme-linked immunosorbent assay (ELISA)

Lentivirus-infected PDX CSC models were used 2 days after selection. Cells were washed with PBS, lysed, and prepared according to the manufacturer’s instructions. Total WNK1 levels were determined using the Human Phospho-WNK1 (T60) & Total WNK1 ELISA Kit (RayBiotech, PEL-WNK1-T60-T). Absorbance was measured on a Victor Nivo multi-mode plate reader.

### Quantitative reverse-transcriptase polymerase chain reaction (qRT-PCR)

RNA was isolated from cells via the RNeasy kit (Qiagen), and concentrations were obtained via a NanoDrop Spectrophotometer. cDNA was subsequently synthesized via qScript cDNA synthesis reagent (Quanta Biosciences) with concentrations measured via the NanoDrop Spectrophotometer. cDNA was diluted to 12.5 ng/µl in accordance with TaqMan (ThermoFisher Scientific) protocols. qPCR was performed on a QuantStudio 3 (Applied Biosystems) thermocycler using TaqMan reactions for Cx43 (*GJA1*; ThermoFisher Scientific, Hs00748445), *WNK1* (ThermoFisher Scientific, Hs01013332), and *C-MYC* (ThermoFisher Scientific, Hs00153408) and TaqMan Universal PCR Master Mix (ThermoFisher Scientific, 4304437). Data were analyzed using the ΔΔCt method and normalized to GAPDH (ThermoFisher Scientific, Hs03929097).

### RNA sequencing

RNA from DI318 CSCs was isolated via RNeasy kit (Qiagen) and submitted to MedGenome for library prep and RNA sequencing using the Illumina TruSeq stranded mRNA kit. Data analysis was performed by MedGenome in accordance with their protocols. RNA-sequencing data for 4121 and 3691 was retrieved from GSE79736^50^. R v4.0.5 was used to create volcano plots and enrichment plots.

Gene set enrichment analysis (GSEA) was individually performed on differentially expressed genes that overlapped between both Cx43 shRNAs (Supplemental Fig. S2) and between both WNK1 shRNAs (Supplemental Fig. S4) compared to non-target via the Molecular Signatures Database (https://www.gsea-msigdb.org/gsea/msigdb/index.jsp), and we focused primarily on transcription factors (TFT:TFT_LEGACY gene sets) to gain mechanistic information. MYC-associated genes were identified from the GSEA datasets: HALLMARK_MYC_TARGETS_V1 (M5926), HALLMARK_MYC_TARGETS_V2 (M5928), COLLER_MYC_TARGETS_UP (M5955), SOUCEK_MYC_TARGETS (M11290), YU_MYC_TARGETS_UP (M1249), and DANG_MYC_TARGETS_UP (M6506).

### Quantification and statistical analysis

All statistical analysis tests were performed in GraphPad Prism 10.2. Information regarding the number of replicates per experiment and specific statistical significance test performed can be found within the individual figure legends. A p-value of < 0.05 was considered statistically significant.

## Results

### Cx43 is expressed by GBM PDX CSCs

Our previous work interrogating the role of connexin proteins in GBM indicated that Cx46 (*GJA3*) is essential for GBM CSC survival, self-renewal, and tumor initiation in a subset of models, with Cx43 primarily expressed in non-stem tumor cells^8,9^. A number of studies using high-passage GBM cells lines and PDX models have suggested that Cx43 (*GJA1*) is also involved in GBM cell proliferation, survival, migration, and therapeutic resistance^18,19,25^. To gain insight into global connexin expression in GBM PDX CSCs, we profiled mRNA expression of all connexins in RNA sequencing datasets from three distinct PDX specimens. Of the 21 human connexin genes, we detected low levels of expression of *GJA3* (Cx46), *GJB1* (Cx32), *GJC1* (Cx45), and *GJC2* (Cx47), with comparatively high expression of *GJA1* (Cx43) (**Fig. 1A**). The relatively high levels of Cx43 and low levels of Cx46 observed here were not consistent with our previous observations and led us to more closely examine levels of Cx43 protein in a larger panel of GBM PDX lines cultured under CSC-promoting conditions. We detected Cx43 protein expression at varying levels in the majority of the examined samples by immunoblot (**Fig. 1B**). Compared to normal human astrocytes (NHAs) as a positive control for Cx43 expression and a panel of non-GBM cancer cell lines (human embryonic kidney line 293T, breast adenocarcinoma cell line MDA-MB-231, cervical cancer line HeLa, and prostate cancer line LNCaP) as a negative control, GBM PDX samples expressed intermediate levels of Cx43 (**Fig. 1C**). Based on our previous work that described an intracellular role for Cx26 in driving triple-negative breast cancer^10,11^, we further interrogated the subcellular localization of Cx43 in GBM PDX models and found that Cx43 was primarily expressed in the membrane fraction, as expected, rather than intracellularly (**Fig. 1D**). Together, these data indicate that Cx43 is expressed at the membrane at detectable levels in GBM PDX CSCs.

**Figure 1.**
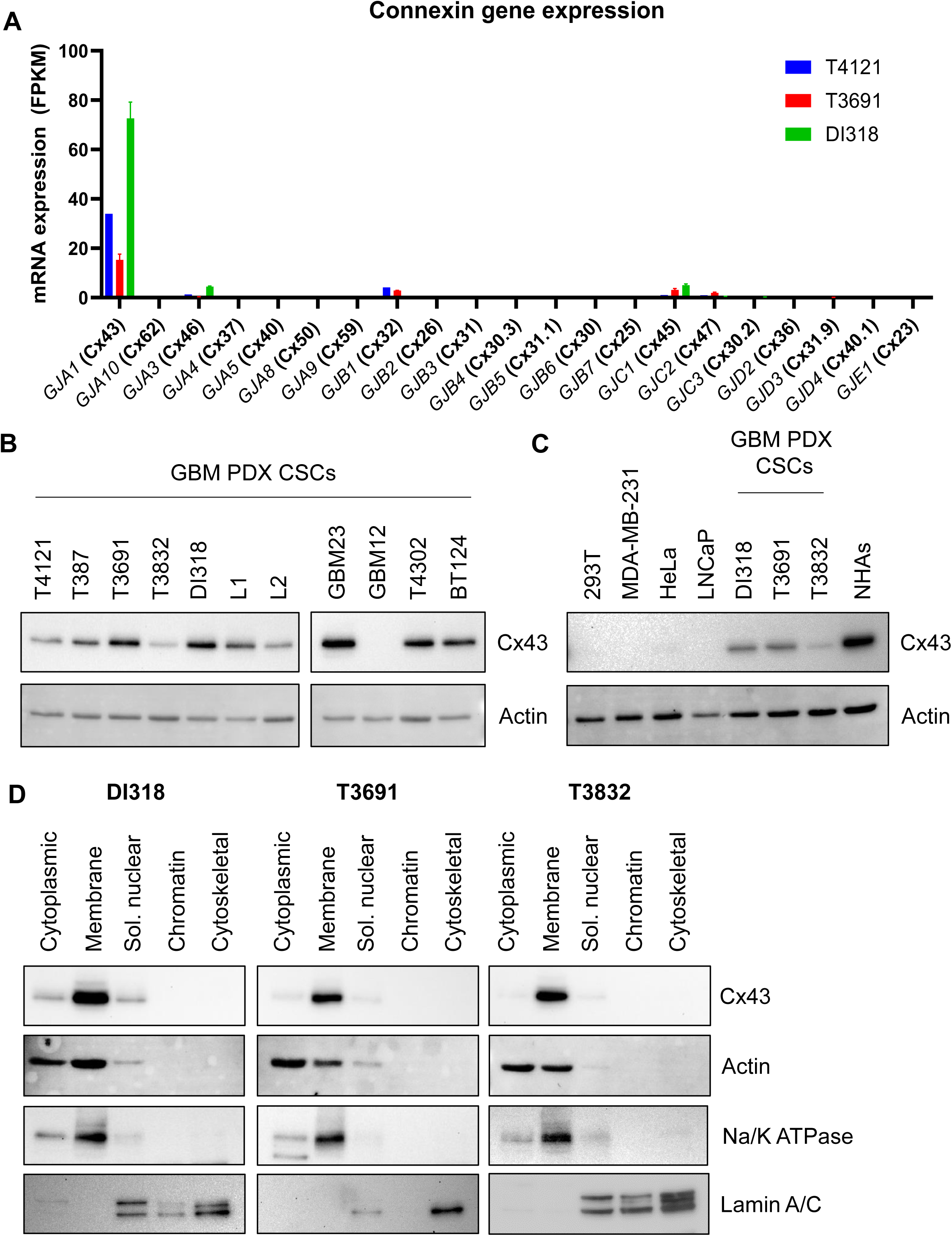
Cx43 is expressed in GBM PDX CSCs. (**A**) Expression of all connexin genes was assessed in GBM PDX CSC models T4121 (GSE79736), T3691 (GSE79736), and DI318 (performed in this study). Expression is shown as fragments per kilobase of transcript per million mapped reads (FPKM). (**B-C**) Baseline levels of Cx43 were examined by immunoblotting in a panel of GBM PDX CSC lines (**B**) and compared those of traditional serum-cultured cell lines and normal human astrocytes (NHAs) (**C**). For B, samples were spread across two gels that were transferred, incubated, and imaged in parallel. Actin was used as a loading control. (**D**) Cx43 is primarily membrane localized in GBM PDX CSCs. Cells were subjected to fractionation as detailed in the Materials and Methods, and immunoblotting of the fractions was performed. The volume used for each fraction was proportional to the number of starting cells. Equal proportions of cytoplasmic, membrane, soluble nuclear, chromatin-bound, and cytoskeletal fractions were loaded. B, C, and D are representative of n=3 independent experiments.

### Cx43 is essential for GBM CSC survival and self-renewal

Based on this observed expression of Cx43 in GBM CSCs, we next interrogated whether Cx43 is important for the function of these cells. Cx43 mRNA was depleted from multiple PDX CSC models using three non-overlapping shRNA constructs (**Fig. 2A, S1A-B**), and we observed a significant decrease in cell number over a span of 5 days (**Fig. 2B, S1C**). This effect was evident even with a relatively modest reduction in Cx43 protein. This reduction in cell number was accompanied by an increase in apoptotic cell death as measured by caspase 3/7 activity (**Fig. 2C, S1D**). We then assayed the critical stem cell property of self-renewal using limiting-dilution analysis and observed a significant decrease in stem cell frequency (**Fig. 2D, S1E**). Together, these results indicate that Cx43 is required for GBM CSC properties including survival and self-renewal and suggest a pro-tumorigenic role for Cx43.

**Figure 2.**
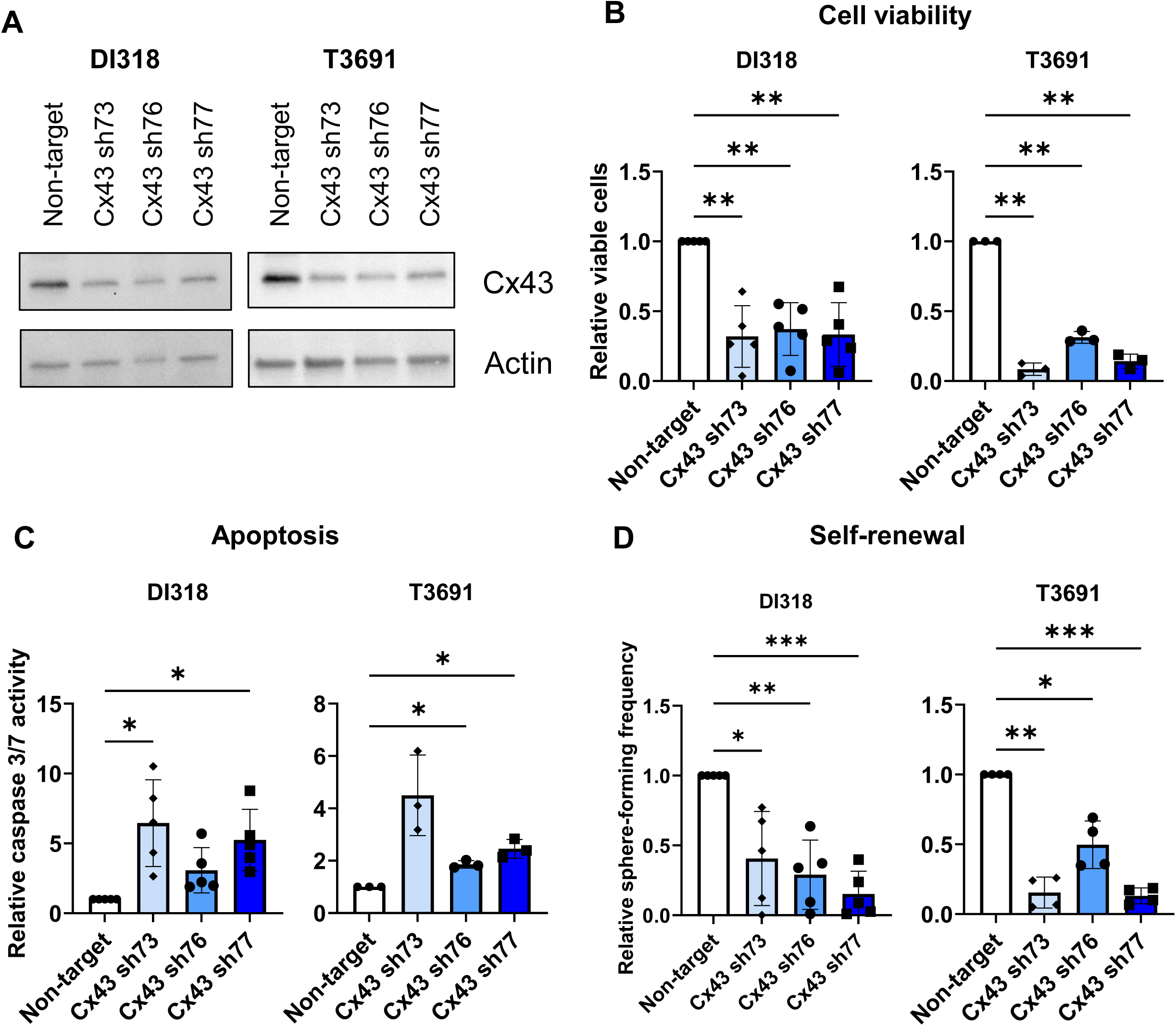
Cx43 is required for GBM PDX CSC survival. DI318 and T3691 PDX CSC models were transduced with three non-overlapping lentiviral shRNA constructs against Cx43 (*GJA1*). (**A**) The degree of knockdown was verified via immunoblotting using an antibody against Cx43. β-actin was used as a loading control. (**B**) Cell viability of DI318 and T3691 CSCs containing shRNAs against Cx43 was measured via CellTiter-Glo on day 5 after plating. Luminescence values were normalized to day 0 values, and the fold change was calculated relative to the non-targeting control. n = 5 (DI318) and n = 3 (T3691) independent experiments, each performed in technical triplicate. (**C**) Cell death of DI318 and T3691 CSCs containing shRNAs against Cx43 was measured via CaspaseGlo 3/7 on day 3 after plating. Luminescence values were normalized to cell number measured via CellTiter Glo at the same time point and then to the non-targeting control. n = 5 (DI318) and n = 3 (T3691) independent experiments, each performed in technical triplicate. (**D**) DI318 and T3691 CSCs containing Cx43 shRNAs were plated in decreasing cell number (20, 10, 5, and 1 cells/well) with 24 replicates per number and evaluated for sphere formation 10-14 days later. n = 5 (DI318) and n = 4 (T3691) independent experiments. The online algorithm outlined in the Methods section was used to calculate stem cell frequency. *p < 0.05, **p < 0.01, ***p < 0.001 by one-way ANOVA analysis with Dunnett’s Multiple Comparison Test.

### Cx43 is required for expression of the c-MYC oncogene

To investigate the mechanistic underpinnings of the requirement for Cx43 in CSCs, we performed RNA sequencing of cells containing shRNA against Cx43. One of the top differences we observed compared to cells expressing a non-target shRNA was a decrease in the levels of the oncogene *c-MYC* (**Fig. 3A, B, Supplemental Fig. S2**) and concomitant decreases in reported MYC-associated genes. MYC is a transcription factor that drives transformation of neural stem cells and is essential for GBM CSC proliferation, survival, self-renewal, and tumor initiation due to its role in cell cycle progression (19020659, 19934320, 18948956, 19150964). We validated the decrease in *MYC* by qRT-PCR in cells expressing Cx43 shRNA (**Fig. 3C**). This change in mRNA levels correlated with a decrease in MYC protein level (**Fig. 3D**). Based on this link between Cx43 and MYC, we next investigated potential signaling intermediates using the Dependency Map (DepMap) portal (https://depmap.org/portal, ^51^). We considered top genes that were found to exhibit co-dependency with *MYC* across a panel of cancer cell lines by either RNAi (DEMETER2 data set^52^; includes the Achilles^51^, DRIVE^53^, and Marcotte^54^ data sets) or CRISPR (DepMap 23Q4)^55^. Only four genes were found in both datasets (**Fig. 3E**), those encoding the MYC dimerization partner MYC-associated factor X (*MAX*), the meiotic protein meiosis regulator for oocyte development (*MIOS*), the mitochondrial protein hydroxysteroid 17-beta dehydrogenase 10 (*HSD17B10*), and the kinase WNK lysine deficient protein kinase 1 (*WNK1*). WNK1 is a member of the WNK family of protein kinases (also including WNK2-4) and canonically regulates ion homeostasis by phosphorylating ion channels^56^. Based on reported roles for WNKs in cancer-associated signaling pathways, including driving expression of *c-MYC*^57–59^, we chose to further investigate the role of WNK1 as a candidate to mediate signaling between Cx43 and *MYC*. We observed a reduction in WNK1 levels by ELISA in GBM CSCs expressing Cx43 shRNA compared to non-target, suggesting a role for Cx43 in WNK1 expression (**Fig. 3F**).

**Figure 3.**
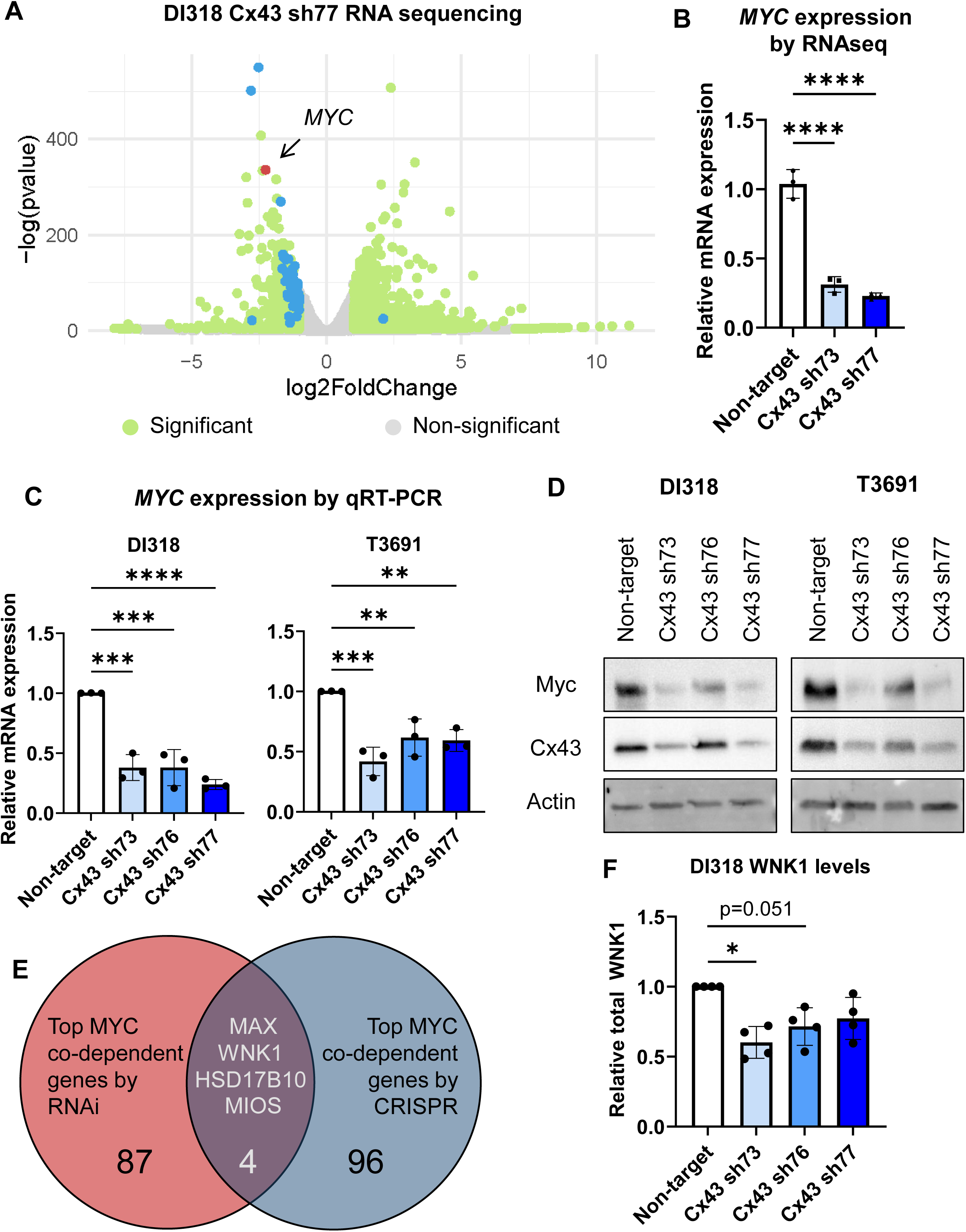
Cx43 is necessary for MYC expression. (**A**) Volcano plot showing differentially expressed genes as determined by RNA sequencing of DI318 PDX CSCs containing Cx43 sh77 compared to non-target. The *MYC* gene is shown in red, with MYC-associated genes shown in blue (see Methods). (**B**) Quantitation of mRNA levels of *MYC* from the RNA sequencing experiment. n=1 experiment with 3 independent replicates. (**C**) *MYC* expression was assessed over three Cx43 shRNA constructs using qRT-PCR in DI318 and T3691 and normalized to GAPDH. n = 3 independent experiments performed in technical triplicate for each model. (**D**) DI318 and T3691 containing Cx43 shRNA were subjected to immunoblot for MYC protein. β-actin was used as a loading control. n ≥ 5 independent replicates. (**E**) The Dependency Map (DepMap) portal was used to identify genes that exhibit co-dependency with *MYC* across a panel of cancer cell lines (see Results section for details). The top approximately 100 hits from both RNAi (DEMETER2 data set) and CRISPR screening (DepMap 23Q4 data set) were selected, and overlapping genes between the two datasets were determined. (**F**) Levels of WNK1 were measured by ELISA in DI318 containing Cx43 shRNA and compared to non-target. n = 4 independent experiments. *p < 0.05, **p < 0.01, ***p < 0.001, **** p < 0.0001 by one-way ANOVA analysis with Dunnett’s Multiple Comparison Test.

### Disruption of WNK1 compromises GBM CSC survival

Based on these data, we interrogated whether WNK1 is important for GBM CSCs. Using three non-overlapping shRNA constructs (**Supplemental Fig. S3A**), we reduced WNK1 levels compared to a non-targeting control (**Fig. 4A, Supplemental Fig. S3B**). These WNK1-depleted cells exhibited a reduction in cell number after 5 days in culture (**Fig. 4B, Supplemental Fig. S3C**) with a concomitant increase in apoptosis as measured by caspase 3/7 activation (**Fig. 4C, Supplemental Fig. S3D**). Self-renewal was also dramatically reduced in cells containing WNK1 shRNA constructs compared to those with a non-targeting control (**Fig. 4D, Supplemental Fig S3E**), suggesting that WNK1 is essential for CSC function. Finally, we implanted CSCs expressing non-target or WNK1 shRNA into the brains of immunocompromised mice and monitored the animals until neurological endpoint. We observed a significant extension in survival of mice with tumors derived from WNK1-depleted cells (**Fig. 4E**), further supporting our observations that WNK1 drives tumor cell growth by GBM CSCs.

**Figure 4.**
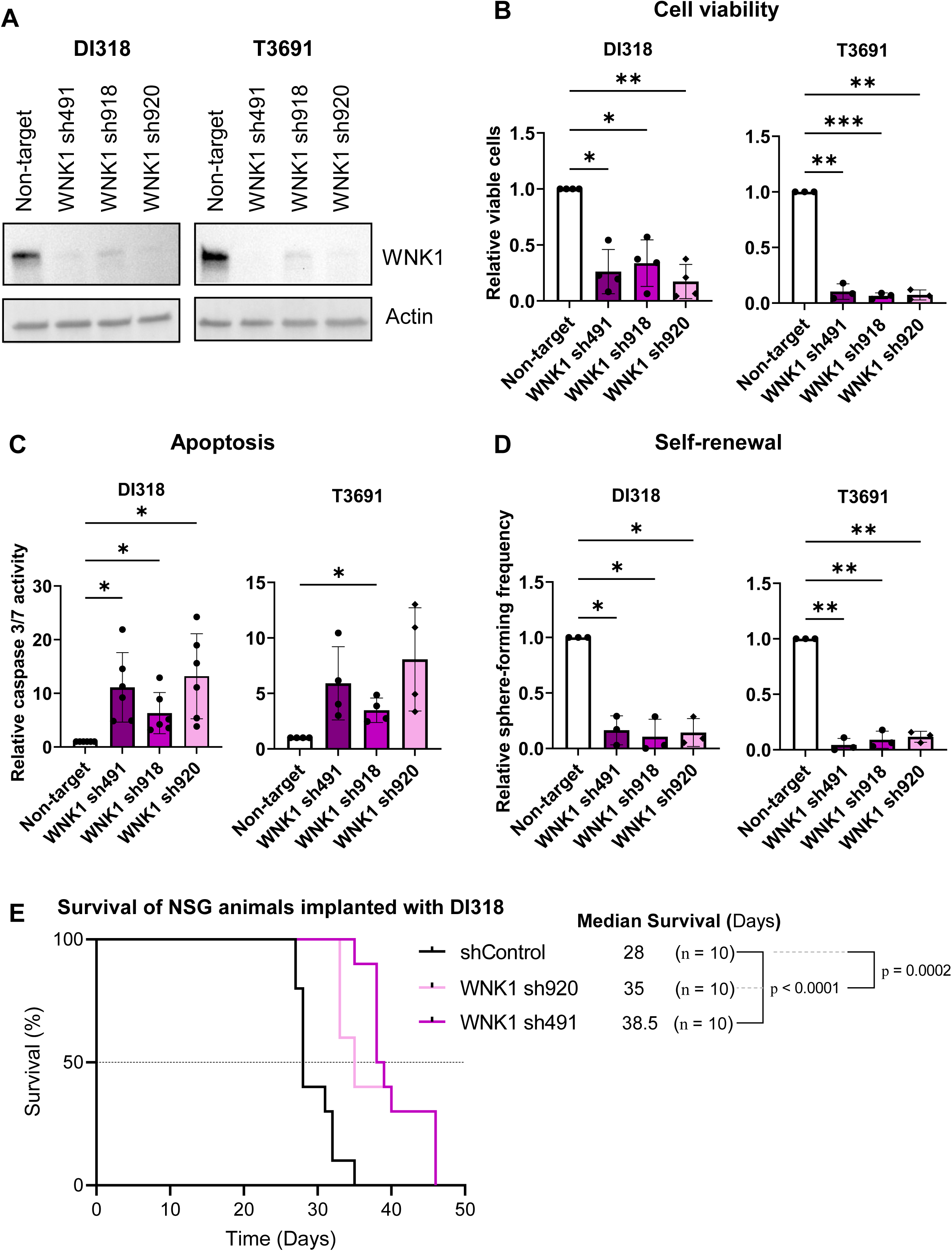
WNK1 is required for CSC viability, survival, and self-renewal. PDX CSC models DI318 and T3691 were transduced with lentivirus encoding three different non-overlapping *WNK1* shRNA constructs. (**A**) Cells were subjected to immunoblotting for WNK1. β-actin was used as a loading control. (**B**) Cell viability of DI318 and T3691 CSCs containing shRNAs against *WNK1* was measured via CellTiter-Glo on day 5 after plating. Luminescence values were normalized to day 0 values, and the fold change was calculated relative to the non-targeting control. n = 4 (DI318) and n = 3 (T3691) independent experiments, each performed in technical triplicate. (**C**) Cell death of DI318 and T3691 CSCs containing shRNAs against *WNK1* was measured via CaspaseGlo 3/7 on day 3 after plating. Luminescence values were normalized to cell number measured via CellTiter Glo at the same time point and then to the non-targeting control. n = 6 (DI318) and n = 4 (T3691) independent experiments, each performed in technical triplicate. (**D**) DI318 and T3691 CSCs containing *WNK1* shRNAs were plated in decreasing cell number (20, 10, 5, and 1 cells/well) with 24 replicates per number and evaluated for sphere formation 10-14 days later. n = 3 independent experiments for both models. The online algorithm outlined in the Methods section was used to calculate stem cell frequency. (**E**) A total of 25,000 DI318 cells containing WNK1 shRNAs was implanted into the brains of immunocompromised NSG mice, and animals were monitored until neurological endpoint. Median survival, animal numbers, and p-values shown on plot; p-value calculated by log-rank test. For panels B-D, *p < 0.05, **p < 0.01, ***p < 0.001 by one-way ANOVA analysis with Dunnett’s Multiple Comparison Test.

### WNK1 controls MYC expression in GBM CSCs

Based on our hypothesis that WNK1 may serve as a signaling intermediate between Cx43 and MYC, we then tested whether WNK1 is necessary for MYC expression. WNK1 has previously been shown to control MYC expression in developing T cells^57^, hepatocellular carcinoma^59^, and multiple myeloma^58^ but not in B cells^60^, but a link in GBM has yet to be determined. To gain a broad understanding of WNK1 mediated signaling changes, we performed RNA sequencing and observed that *MYC* and MYC targets were significantly downregulated in GBM CSCs containing WNK1 shRNA compared to a non-targeting control (**Fig. 5A, B, Supplemental Fig. S4A**). We further confirmed this result by qRT-PCR and observed a significant reduction in *MYC* mRNA levels in the presence of WNK1 shRNA (**Fig. 5C**). MYC protein showed a similar trend, with decreases in the presence of WNK1 shRNAs compared to a non-targeting sequence (**Fig. 5D, Supplemental Fig. S4B**). As WNK2 has tumor-suppressive effects in GBM cells^61,62^, we utilized RNA sequencing data to determine whether WNK1 depletion might act by increasing WNK2 levels. Importantly, RNA sequencing revealed that expression of *WNK2*, *WNK3*, *and WNK4* was very low in GBM CSCs and did not increase with Cx43 or WNK1 shRNAs (**Supplemental Fig. S4C**), suggesting that other WNK proteins do not compensate at least at the transcriptional level upon Cx43 or WNK1 depletion. Based on our observations that depletion of WNK1 phenocopies reduction of Cx43 and Cx43 reduces levels of WNK1, together, our work suggests the presence of a signaling axis containing Cx43, WNK1, and MYC that is essential for GBM CSCs.

**Figure 5.**
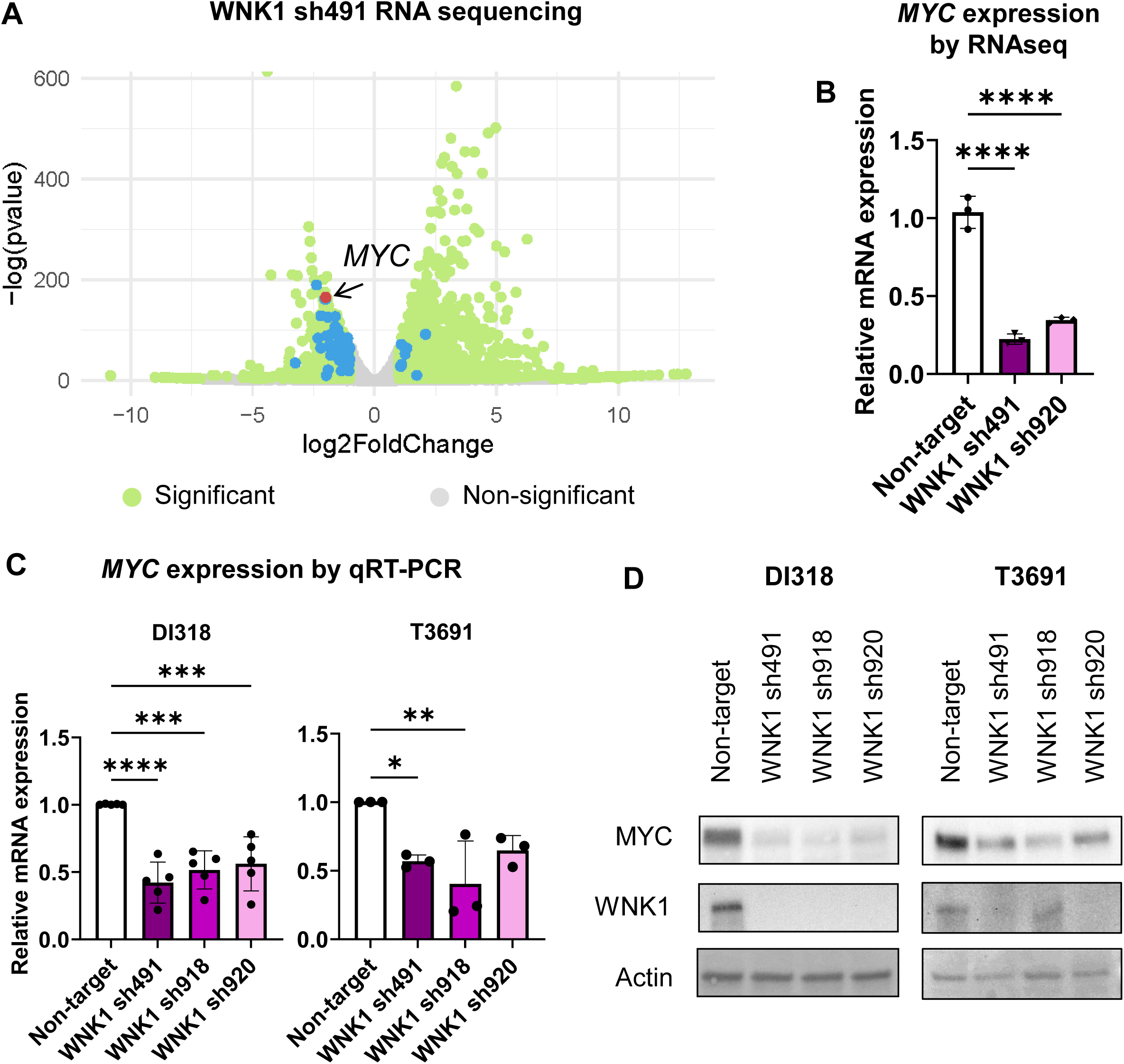
WNK1 is upstream of MYC in GBM CSCs. (**A**) Volcano plot of significantly differentially expressed genes from RNA sequencing of DI318 cells containing *WNK1* sh491. *MYC* is shown in red, with MYC-associated genes in blue. (**B**) Quantitation of *MYC* mRNA level by RNAseq in cells containing *WNK1* shRNA compared to non-targeting control. n = 1 experiment of three independent replicates. (**C**) *MYC* expression was assessed over three *WNK1* shRNA constructs using qRT-PCR in DI318 and T3691 and normalized to GAPDH. n = 5 (DI318) and n = 3 (T3691) independent experiments performed in technical triplicate for each model. (**D**) DI318 and T3691 containing *WNK1* shRNA were subjected to immunoblotting for MYC protein. β-actin was used as a loading control. n ≥ 5 independent replicates. *p < 0.05, **p < 0.01, ***p < 0.001 by one-way ANOVA analysis with Dunnett’s Multiple Comparison Test.

## Discussion

While our understanding of the tumor-suppressive and tumor-promoting roles of connexins continues to evolve, our observations demonstrate a new pro-tumorigenic signaling axis in GBM CSCs linking Cx43 to c-MYC via WNK1. Our previous work showed an essential role for GJIC through Cx46 in GBM CSCs from a subset of patient samples^8,9^. However, this work was unable to directly compare levels of Cx46 to those of other connexins. Here, using sequencing approaches, we show that each of the GBM CSC samples assessed expressed higher levels of Cx43 compared to all other connexins at the mRNA level. The question remains as to whether these patient samples exhibit distinct dependencies on an individual connexin or whether cells that require Cx46 may also require Cx43. The levels of Cx43 we detected in GBM cells were higher than those of traditional, serum-cultured cancer cell lines but lower than those observed in normal astrocytes. The low to undetectable levels in breast cancer, cervical cancer, and prostate cancer cell lines are consistent with the traditional hypothesis that Cx43 is tumor suppressive and is lost upon transformation. However, the comparatively robust expression of Cx43 in GBM CSCs contrasts with that hypothesis and lends further support to the idea that context is critical for understanding the role of connexins and gap junctions, with varying expression, dependency, and molecular function depending on the situation. The mechanisms underlying this difference in expression remain unexplored; it is possible that Cx43 levels are regulated by culture conditions, as we observed low expression with the presence of serum. However, our normal astrocytes with high Cx43 were also grown in the presence of serum, although a reduced amount. Alternatively, our interrogation of the CSC fraction of GBM rather than the bulk tumor may have enriched for Cx43 expression; the low frequency of CSCs within the tumor bulk may in fact have contributed to the observation in historical studies of reduced levels of Cx43 within GBM. It is also possible that GBM may simply differ in its underlying biology compared to non-neural tumors, or the long-term culture of the traditional cell lines has led to reduced Cx43 levels. This question of how our GBM models express notable levels of Cx43 protein warrants additional study.

Here, we present data indicating a pro-tumorigenic role for Cx43 in GBM CSCs. Mechanistically, we observed that expression of both transcript and protein levels of the oncogene c-MYC relies on the presence of Cx43 and WNK1. To our knowledge, this is the first time a link between Cx43 and MYC or between Cx43 and WNK1 has been reported. c-MYC is a critical regulator of cell cycle progression and apoptosis that is frequently overexpressed in GBM^63^. Previous work from our group and others showed this essential role for MYC in GBM CSCs: MYC is necessary for cell survival, cell-cycle progression, self-renewal, and tumor formation^64,65^. Based on these data, we believe that the loss of MYC is likely sufficient to drive the reduced survival we observed upon Cx43 depletion. However, Cx43 likely affects numerous other cellular processes, as supported by our RNA sequencing data, some of which may also be essential for cell survival.

While WNK1 has previously been identified upstream of MYC expression in hepatocellular carcinoma^59^, multiple myeloma^58^, and developing T cells^57^, this is the first report linking WNK1 and MYC in GBM, indicating that WNK1 is an important and understudied player in GBM biology. Our data show that WNK1 supports tumor cell growth, and this is supported by the small amount of previous work on WNK1 in GBM. Inhibition of WNK1 immediate downstream substrates, STE20/SPS1-related proline/alanine-rich kinase (SPAK) and oxidative stress-responsive kinase 1 (OSR1), reduced glioblastoma proliferation, migration, and tumor growth^66^; however this study did not test for any kinase- or SPAK-independent roles for WNK1. Another study also detected reduced glioma cell migration with WNK1 shRNA, likely due to reduced phosphorylation of Na+/K+/2Cl− cotransporter isoform 1 (NKCC1), but did not assess a role in cellular proliferation, stem cell functions, or tumor growth^67^. While we were unable to detect mRNA expression of any other WNK family member in our cells (**Supplemental Fig. S4C**), WNK3 has also been shown to promote GBM cell migration^68,69^. Interestingly, while WNK1 and WNK3 appear to be tumor promoting, WNK2 has been shown to suppress glioma cell migration^70^, expression of matrix metalloproteinase 2 (MMP2)^62^, and activation of c-Jun N-terminal kinase (JNK)^62^, suggesting that it instead has tumor-suppressive functions. Furthermore, WNK1 has a MYC response element in its promoter, suggesting that the relationship between these two proteins may be more complex than currently understood. Additional work is needed to fully understand the roles of the WNK family members in tumor cell properties such as growth, migration, and tumor formation and their relationship to MYC expression.

The Cx43-WNK1-MYC signaling axis may present opportunities for therapeutic development. Connexins have historically been difficult to specifically target due to the high degree of sequence homology among family members. However, several Cx43-specific targeting strategies have been developed in recent years. αCT1 is a peptide mimetic of the region of the Cx43 C-terminal intracellular tail that binds to zonula occludens 1 (ZO-1), which regulates the balance between docked and undocked connexon hemichannels at the membrane^71,72^. Treatment of cells with αCT1 increases gap junction incorporation of Cx43 while reducing hemichannel activity^71,72^ and has been tested clinically to increase wound healing^73^. However, as our data support a pro-tumorigenic role for Cx43, increasing the GJIC of Cx43 is likely to be detrimental rather than beneficial to patients. An additional peptide mimetic of Cx43, TAT-Cx43_266-283_, has been developed that inhibits the activity of SRC^36,39,40^. This peptide is based on the observation that the presence of Cx43 suppresses SRC activity and again mimics the presence of Cx43 rather than reducing its function. Thus, the available Cx43 peptide treatments are not likely to be useful for reducing tumor growth in GBM cells expressing Cx43; however, these types of therapeutic strategies do suggest that it may be possible to develop specific therapies targeting Cx43 in the future. WNK1 and MYC targeting strategies have also encountered roadblocks. In addition to cancers, therapies to target WNK1 have been of interest to treat Gordon’s syndrome (also called pseudohypoaldosteronism type II (PHA2)), a type of familial hypertension caused by WNK mutations. However, WNK inhibitors lack specificity for a single WNK family member, leading to undesirable effects, and development of the WNK inhibitor WNK463 was halted due to unexpected side effects in mice^74^. MYC inhibitors have also been highly sought after for use in cancer treatment, but targeting MYC has thus far been challenging due to its highly disordered regions and lack of an enzymatic active site^75^. While several strategies to target MYC are currently in clinical trials, including the MYC-MAX interaction inhibitor omomyc and a number of indirect inhibitors that target upstream proteins, none have yet been approved for use in the clinic^75^. The increased understanding of additional signaling nodes surrounding poorly targetable proteins such as Cx43, WNK1, and MYC may provide more amenable targets for therapeutic development. This strategy is particularly relevant for proteins such as Cx43 that have multifaceted roles in tumor cells.

Together, our work demonstrates a critical role for Cx43 and WNK1 in GBM CSCs for driving MYC expression, leading to cell survival, self-renewal, and tumor formation. However, there are several limitations to our study. While we thoroughly investigated three PDX GBM models, the future investigation of additional models will be beneficial, particularly due to the high degree of heterogeneity present in GBM. In addition, with recent work suggesting sex differences may impact GBM cell phenotypes, including cell growth^76^ and immune responses^77,78^, a more detailed assessment of this signaling axis with equal representation of male and female models would be useful. Our studies focused on the cell signaling network starting from Cx43, but it is unclear which function(s) of Cx43 (GJIC, hemichannel, or protein-protein interactions) is responsible for the initiation of this signaling network and whether there are any points of feedback. These are key questions to be addressed in future studies, along with whether the WNK1/c-MYC signaling axis can be initiated by any other connexin. Finally, future studies would benefit from focusing on untangling the role WNK1 kinase activity in the phenotypes observed here, as many additional components of this pathway remain to be discovered.

## Acknowledgements

We thank the members of the Lathia laboratory for insightful discussion and constructive comments on the manuscript and Dr. Reza Khatib for his inspiration and support of our work. We greatly appreciate the illustrative work of Ms. Amanda Mendelsohn from the Center for Medical Art and Photography at the Cleveland Clinic. This work was funded by support from the NIH (R01 NS089641), the Cleveland Clinic VeloSano Bike Race, the American Cancer Society (Research Scholar Grant), the Case Comprehensive Cancer Center, and the Cleveland Clinic/Lerner Research Institute to JDL. DJS was supported by an ABTA Discovery Award (DG2300058) and the VeloSano Bike Race.

## Author contributions

Conceptualization: EEM, NH, CGH, JDL; methodology: EEM, NH; Formal analysis: EEM, NH, EH, DJS, CGH; Investigation: EEM, NH, EH, APJ, KEK, SZW, DJS, CGH; Resources: JDL; Writing - Original Draft: EEM, NH; Writing - Review & Editing: EEM, NH, JDL; Visualization: EEM, EH; Supervision: EEM, JDL; Funding Acquisition: EEM, JDL; approval of the final version: all authors.

## Declaration of interests

The authors declare no competing interests.

## Notes

### Competing Interest Statement

The authors have declared no competing interest.

## References

1. Stupp R, Mason WP, van den Bent MJ, Weller M, Fisher B, Taphoorn MJ, Belanger K, Brandes AA, Marosi C, Bogdahn U, et al. Radiotherapy plus concomitant and adjuvant temozolomide for glioblastoma. N Engl J Med. 2005;352:987–996. doi: 10.1056/NEJMoa043330

2. Aasen T, Mesnil M, Naus CC, Lampe PD, Laird DW. Gap junctions and cancer: communicating for 50 years. Nat Rev Cancer. 2016;16:775–788. doi: 10.1038/nrc.2016.105

3. Mulkearns-Hubert EE, Reizes O, Lathia JD. Connexins in Cancer: Jekyll or Hyde? Biomolecules. 2020;10. doi: 10.3390/biom10121654

4. Saez JC, Leybaert L. Hunting for connexin hemichannels. FEBS Lett. 2014;588:1205–1211. doi: 10.1016/j.febslet.2014.03.004

5. Loewenstein WR, Kanno Y. Intercellular communication and the control of tissue growth: lack of communication between cancer cells. Nature. 1966;209:1248–1249. doi: 10.1038/2091248a0

6. Sin WC, Crespin S, Mesnil M. Opposing roles of connexin43 in glioma progression. Biochim Biophys Acta. 2012;1818:2058–2067. doi: 10.1016/j.bbamem.2011.10.022

7. Mesnil M, Crespin S, Avanzo JL, Zaidan-Dagli ML. Defective gap junctional intercellular communication in the carcinogenic process. Biochim Biophys Acta. 2005;1719:125–145. doi: 10.1016/j.bbamem.2005.11.004

8. Hitomi M, Deleyrolle LP, Mulkearns-Hubert EE, Jarrar A, Li M, Sinyuk M, Otvos B, Brunet S, Flavahan WA, Hubert CG, et al. Differential connexin function enhances self-renewal in glioblastoma. Cell reports. 2015;11:1031–1042. doi: 10.1016/j.celrep.2015.04.021

9. Mulkearns-Hubert EE, Torre-Healy LA, Silver DJ, Eurich JT, Bayik D, Serbinowski E, Hitomi M, Zhou J, Przychodzen B, Zhang R, et al. Development of a Cx46 Targeting Strategy for Cancer Stem Cells. Cell reports. 2019;27:1062–1072 e1065. doi: 10.1016/j.celrep.2019.03.079

10. Thiagarajan PS, Sinyuk M, Turaga SM, Mulkearns-Hubert EE, Hale JS, Rao V, Demelash A, Saygin C, China A, Alban TJ, et al. Cx26 drives self-renewal in triple-negative breast cancer via interaction with NANOG and focal adhesion kinase. Nat Commun. 2018;9:578. doi: 10.1038/s41467-018-02938-1

11. Mulkearns-Hubert EE, Esakov Rhoades E, Ben-Salem S, Bharti R, Hajdari N, Johnson S, Myers A, Smith IN, Bandyopadhyay S, Eng C, et al. Targeting NANOG and FAK via Cx26-derived Cell-penetrating Peptides in Triple-negative Breast Cancer. Mol Cancer Ther. 2024;23:56–67. doi: 10.1158/1535-7163.MCT-21-0783

12. Shinoura N, Chen L, Wani MA, Kim YG, Larson JJ, Warnick RE, Simon M, Menon AG, Bi WL, Stambrook PJ. Protein and messenger RNA expression of connexin43 in astrocytomas: implications in brain tumor gene therapy. Journal of neurosurgery. 1996;84:839–845; discussion 846. doi: 10.3171/jns.1996.84.5.0839

13. Estin D, Li M, Spray D, Wu JK. Connexins are expressed in primary brain tumors and enhance the bystander effect in gene therapy. Neurosurgery. 1999;44:361–368; discussion 368-369. doi: 10.1097/00006123-199902000-00068

14. Aronica E, Gorter JA, Jansen GH, Leenstra S, Yankaya B, Troost D. Expression of connexin 43 and connexin 32 gap-junction proteins in epilepsy-associated brain tumors and in the perilesional epileptic cortex. Acta Neuropathol. 2001;101:449–459. doi: 10.1007/s004010000305

15. Huang RP, Hossain MZ, Sehgal A, Boynton AL. Reduced connexin43 expression in high-grade human brain glioma cells. J Surg Oncol. 1999;70:21–24. doi: 10.1002/(sici)1096-9098(199901)70:1<21::aid-jso4>3.0.co;2-0

16. Soroceanu L, Manning TJ, Jr., Sontheimer H. Reduced expression of connexin-43 and functional gap junction coupling in human gliomas. Glia. 2001;33:107–117. doi: 10.1002/1098-1136(200102)33:2<107::aid-glia1010>3.0.co;2-4

17. Gielen PR, Aftab Q, Ma N, Chen VC, Hong X, Lozinsky S, Naus CC, Sin WC. Connexin43 confers Temozolomide resistance in human glioma cells by modulating the mitochondrial apoptosis pathway. Neuropharmacology. 2013;75:539–548. doi: 10.1016/j.neuropharm.2013.05.002

18. Murphy SF, Varghese RT, Lamouille S, Guo S, Pridham KJ, Kanabur P, Osimani AM, Sharma S, Jourdan J, Rodgers CM, et al. Connexin 43 Inhibition Sensitizes Chemoresistant Glioblastoma Cells to Temozolomide. Cancer Res. 2016;76:139–149. doi: 10.1158/0008-5472.CAN-15-1286

19. Pridham KJ, Shah F, Hutchings KR, Sheng KL, Guo S, Liu M, Kanabur P, Lamouille S, Lewis G, Morales M, et al. Connexin 43 confers chemoresistance through activating PI3K. Oncogenesis. 2022;11:2. doi: 10.1038/s41389-022-00378-7

20. Che J, DePalma TJ, Sivakumar H, Mezache LS, Tallman MM, Venere M, Swindle-Reilly K, Veeraraghavan R, Skardal A. alphaCT1 peptide sensitizes glioma cells to temozolomide in a glioblastoma organoid platform. Biotechnol Bioeng. 2023;120:1108–1119. doi: 10.1002/bit.28313

21. Grek CL, Sheng Z, Naus CC, Sin WC, Gourdie RG, Ghatnekar GG. Novel approach to temozolomide resistance in malignant glioma: connexin43-directed therapeutics. Curr Opin Pharmacol. 2018;41:79–88. doi: 10.1016/j.coph.2018.05.002

22. Chen W, Wang D, Du X, He Y, Chen S, Shao Q, Ma C, Huang B, Chen A, Zhao P, et al. Glioma cells escaped from cytotoxicity of temozolomide and vincristine by communicating with human astrocytes. Med Oncol. 2015;32:43. doi: 10.1007/s12032-015-0487-0

23. Yusubalieva GM, Baklaushev VP, Gurina OI, Zorkina YA, Gubskii IL, Kobyakov GL, Golanov AV, Goryainov SA, Gorlachev GE, Konovalov AN, et al. Treatment of poorly differentiated glioma using a combination of monoclonal antibodies to extracellular connexin-43 fragment, temozolomide, and radiotherapy. Bull Exp Biol Med. 2014;157:510–515. doi: 10.1007/s10517-014-2603-0

24. Munoz JL, Rodriguez-Cruz V, Greco SJ, Ramkissoon SH, Ligon KL, Rameshwar P. Temozolomide resistance in glioblastoma cells occurs partly through epidermal growth factor receptor-mediated induction of connexin 43. Cell Death Dis. 2014;5:e1145. doi: 10.1038/cddis.2014.111

25. Wang GG, Wang Y, Wang SL, Zhu LC. Down-regulation of CX43 expression by miR-1 inhibits the proliferation and invasion of glioma cells. Transl Cancer Res. 2022;11:4126–4136. doi: 10.21037/tcr-22-2318

26. Krigers A, Moser P, Fritsch H, Demetz M, Kerschbaumer J, Brawanski KR, Thome C, Freyschlag CF. The relationship between connexin-43 expression and Ki67 in non-glial central nervous system tumors. Int J Biol Markers. 2023;38:46–52. doi: 10.1177/03936155221143138

27. Osswald M, Jung E, Sahm F, Solecki G, Venkataramani V, Blaes J, Weil S, Horstmann H, Wiestler B, Syed M, et al. Brain tumour cells interconnect to a functional and resistant network. Nature. 2015;528:93–98. doi: 10.1038/nature16071

28. McCutcheon S, Spray DC. Glioblastoma-Astrocyte Connexin 43 Gap Junctions Promote Tumor Invasion. Mol Cancer Res. 2022;20:319–331. doi: 10.1158/1541-7786.MCR-21-0199

29. Lin JH, Takano T, Cotrina ML, Arcuino G, Kang J, Liu S, Gao Q, Jiang L, Li F, Lichtenberg-Frate H, et al. Connexin 43 enhances the adhesivity and mediates the invasion of malignant glioma cells. J Neurosci. 2002;22:4302–4311. doi: 10.1523/JNEUROSCI.22-11-04302.2002

30. Zhu D, Caveney S, Kidder GM, Naus CC. Transfection of C6 glioma cells with connexin 43 cDNA: analysis of expression, intercellular coupling, and cell proliferation. Proceedings of the National Academy of Sciences of the United States of America. 1991;88:1883–1887. doi: 10.1073/pnas.88.5.1883

31. Huang RP, Fan Y, Hossain MZ, Peng A, Zeng ZL, Boynton AL. Reversion of the neoplastic phenotype of human glioblastoma cells by connexin 43 (cx43). Cancer Res. 1998;58:5089–5096.

32. Xia Z, Pu P, Huang Q. [Effect of transfected Cx43 gene on the gap junction intercellular communication and the human glioma cells proliferation]. Zhonghua Zhong Liu Za Zhi. 2001;23:465–468.

33. Jaraiz-Rodriguez M, Tabernero MD, Gonzalez-Tablas M, Otero A, Orfao A, Medina JM, Tabernero A. A Short Region of Connexin43 Reduces Human Glioma Stem Cell Migration, Invasion, and Survival through Src, PTEN, and FAK. Stem Cell Reports. 2017;9:451–463. doi: 10.1016/j.stemcr.2017.06.007

34. Strale PO, Clarhaut J, Lamiche C, Cronier L, Mesnil M, Defamie N. Down-regulation of Connexin43 expression reveals the involvement of caveolin-1 containing lipid rafts in human U251 glioblastoma cell invasion. Mol Carcinog. 2012;51:845–860. doi: 10.1002/mc.20853

35. Lu J, Yu M, Lin Z, Lue S, Zhang H, Zhao H, Xu Y, Liu H. Effects of Connexin43 Overexpression on U251 Cell Growth, Migration, and Apoptosis. Med Sci Monit. 2017;23:2917–2923. doi: 10.12659/msm.905130

36. Gonzalez-Sanchez A, Jaraiz-Rodriguez M, Dominguez-Prieto M, Herrero-Gonzalez S, Medina JM, Tabernero A. Connexin43 recruits PTEN and Csk to inhibit c-Src activity in glioma cells and astrocytes. Oncotarget. 2016;7:49819–49833. doi: 10.18632/oncotarget.10454

37. McDonough WS, Johansson A, Joffee H, Giese A, Berens ME. Gap junction intercellular communication in gliomas is inversely related to cell motility. Int J Dev Neurosci. 1999;17:601–611. doi: 10.1016/s0736-5748(99)00024-6

38. Lee J, Kotliarova S, Kotliarov Y, Li A, Su Q, Donin NM, Pastorino S, Purow BW, Christopher N, Zhang W, et al. Tumor stem cells derived from glioblastomas cultured in bFGF and EGF more closely mirror the phenotype and genotype of primary tumors than do serum-cultured cell lines. Cancer cell. 2006;9:391–403. doi: 10.1016/j.ccr.2006.03.030

39. Gangoso E, Thirant C, Chneiweiss H, Medina JM, Tabernero A. A cell-penetrating peptide based on the interaction between c-Src and connexin43 reverses glioma stem cell phenotype. Cell Death Dis. 2014;5:e1023. doi: 10.1038/cddis.2013.560

40. Jaraiz-Rodriguez M, Talaveron R, Garcia-Vicente L, Pelaz SG, Dominguez-Prieto M, Alvarez-Vazquez A, Flores-Hernandez R, Sin WC, Bechberger J, Medina JM, et al. Connexin43 peptide, TAT-Cx43266-283, selectively targets glioma cells, impairs malignant growth, and enhances survival in mouse models in vivo. Neuro-oncology. 2020;22:493–504. doi: 10.1093/neuonc/noz243

41. Alvarado AG, Thiagarajan PS, Mulkearns-Hubert EE, Silver DJ, Hale JS, Alban TJ, Turaga SM, Jarrar A, Reizes O, Longworth MS, et al. Glioblastoma Cancer Stem Cells Evade Innate Immune Suppression of Self-Renewal through Reduced TLR4 Expression. Cell stem cell. 2017;20:450–461 e454. doi: 10.1016/j.stem.2016.12.001

42. Alvarado AG, Turaga SM, Sathyan P, Mulkearns-Hubert EE, Otvos B, Silver DJ, Hale JS, Flavahan WA, Zinn PO, Sinyuk M, et al. Coordination of self-renewal in glioblastoma by integration of adhesion and microRNA signaling. Neuro-oncology. 2016;18:656–666. doi: 10.1093/neuonc/nov196

43. Guryanova OA, Wu Q, Cheng L, Lathia JD, Huang Z, Yang J, MacSwords J, Eyler CE, McLendon RE, Heddleston JM, et al. Nonreceptor tyrosine kinase BMX maintains self-renewal and tumorigenic potential of glioblastoma stem cells by activating STAT3. Cancer cell. 2011;19:498–511. doi: 10.1016/j.ccr.2011.03.004

44. Mitchell K, Sprowls SA, Arora S, Shakya S, Silver DJ, Goins CM, Wallace L, Roversi G, Schafer RE, Kay K, et al. WDR5 represents a therapeutically exploitable target for cancer stem cells in glioblastoma. Genes Dev. 2023;37:86–102. doi: 10.1101/gad.349803.122

45. Cusulin C, Chesnelong C, Bose P, Bilenky M, Kopciuk K, Chan JA, Cairncross JG, Jones SJ, Marra MA, Luchman HA, et al. Precursor States of Brain Tumor Initiating Cell Lines Are Predictive of Survival in Xenografts and Associated with Glioblastoma Subtypes. Stem Cell Reports. 2015;5:1–9. doi: 10.1016/j.stemcr.2015.05.010

46. Siebzehnrubl FA, Silver DJ, Tugertimur B, Deleyrolle LP, Siebzehnrubl D, Sarkisian MR, Devers KG, Yachnis AT, Kupper MD, Neal D, et al. The ZEB1 pathway links glioblastoma initiation, invasion and chemoresistance. EMBO Mol Med. 2013;5:1196–1212. doi: 10.1002/emmm.201302827

47. Deleyrolle LP, Harding A, Cato K, Siebzehnrubl FA, Rahman M, Azari H, Olson S, Gabrielli B, Osborne G, Vescovi A, et al. Evidence for label-retaining tumour-initiating cells in human glioblastoma. Brain. 2011;134:1331–1343. doi: 10.1093/brain/awr081

48. Silver DJ, Roversi GA, Bithi N, Wang SZ, Troike KM, Neumann CK, Ahuja GK, Reizes O, Brown JM, Hine C, et al. Severe consequences of a high-lipid diet include hydrogen sulfide dysfunction and enhanced aggression in glioblastoma. J Clin Invest. 2021;131. doi: 10.1172/JCI138276

49. Hu Y, Smyth GK. ELDA: extreme limiting dilution analysis for comparing depleted and enriched populations in stem cell and other assays. J Immunol Methods. 2009;347:70–78. doi: 10.1016/j.jim.2009.06.008

50. Xie Q, Wu Q, Kim L, Miller TE, Liau BB, Mack SC, Yang K, Factor DC, Fang X, Huang Z, et al. RBPJ maintains brain tumor-initiating cells through CDK9-mediated transcriptional elongation. J Clin Invest. 2016;126:2757–2772. doi: 10.1172/JCI86114

51. Tsherniak A, Vazquez F, Montgomery PG, Weir BA, Kryukov G, Cowley GS, Gill S, Harrington WF, Pantel S, Krill-Burger JM, et al. Defining a Cancer Dependency Map. Cell. 2017;170:564–576 e516. doi: 10.1016/j.cell.2017.06.010

52. McFarland JM, Ho ZV, Kugener G, Dempster JM, Montgomery PG, Bryan JG, Krill-Burger JM, Green TM, Vazquez F, Boehm JS, et al. Improved estimation of cancer dependencies from large-scale RNAi screens using model-based normalization and data integration. Nat Commun. 2018;9:4610. doi: 10.1038/s41467-018-06916-5

53. McDonald ER, 3rd, de Weck A, Schlabach MR, Billy E, Mavrakis KJ, Hoffman GR, Belur D, Castelletti D, Frias E, Gampa K, et al. Project DRIVE: A Compendium of Cancer Dependencies and Synthetic Lethal Relationships Uncovered by Large-Scale, Deep RNAi Screening. Cell. 2017;170:577–592 e510. doi: 10.1016/j.cell.2017.07.005

54. Marcotte R, Sayad A, Brown KR, Sanchez-Garcia F, Reimand J, Haider M, Virtanen C, Bradner JE, Bader GD, Mills GB, et al. Functional Genomic Landscape of Human Breast Cancer Drivers, Vulnerabilities, and Resistance. Cell. 2016;164:293–309. doi: 10.1016/j.cell.2015.11.062

55. DepMap B [database online]. Figshare+: 2023.

56. Vitari AC, Deak M, Morrice NA, Alessi DR. The WNK1 and WNK4 protein kinases that are mutated in Gordon’s hypertension syndrome phosphorylate and activate SPAK and OSR1 protein kinases. Biochem J. 2005;391:17–24. doi: 10.1042/BJ20051180

57. Kochl R, Vanes L, Llorian Sopena M, Chakravarty P, Hartweger H, Fountain K, White A, Cowan J, Anderson G, Tybulewicz VL. Critical role of WNK1 in MYC-dependent early mouse thymocyte development. Elife. 2020;9. doi: 10.7554/eLife.56934

58. Ye T, Mishra AK, Banday S, Li R, Hu K, Coleman MM, Shan Y, Chowdhury SR, Zhou L, Pak ML, et al. Identification of WNK1 as a therapeutic target to suppress IgH/MYC expression in multiple myeloma. Cell reports. 2024;43:114211. doi: 10.1016/j.celrep.2024.114211

59. Teng F, Guo M, Liu F, Wang C, Dong J, Zhang L, Zou Y, Chen R, Sun K, Fu H, et al. Treatment with an SLC12A1 antagonist inhibits tumorigenesis in a subset of hepatocellular carcinomas. Oncotarget. 2016;7:53571–53582. doi: 10.18632/oncotarget.10670

60. Hayward DA, Vanes L, Wissmann S, Sivapatham S, Hartweger H, O’May JB, de Boer LL, Mitter R, Kochl R, Stein JV, et al. B cell-intrinsic requirement for WNK1 kinase in antibody responses in mice. J Exp Med. 2023;220. doi: 10.1084/jem.20211827

61. Hong C, Moorefield KS, Jun P, Aldape KD, Kharbanda S, Phillips HS, Costello JF. Epigenome scans and cancer genome sequencing converge on WNK2, a kinase-independent suppressor of cell growth. Proceedings of the National Academy of Sciences of the United States of America. 2007;104:10974–10979. doi: 10.1073/pnas.0700683104

62. Costa AM, Pinto F, Martinho O, Oliveira MJ, Jordan P, Reis RM. Silencing of the tumor suppressor gene WNK2 is associated with upregulation of MMP2 and JNK in gliomas. Oncotarget. 2015;6:1422–1434. doi: 10.18632/oncotarget.2805

63. Herms JW, von Loewenich FD, Behnke J, Markakis E, Kretzschmar HA. c-myc oncogene family expression in glioblastoma and survival. Surg Neurol. 1999;51:536–542. doi: 10.1016/s0090-3019(98)00028-7

64. Wang J, Wang H, Li Z, Wu Q, Lathia JD, McLendon RE, Hjelmeland AB, Rich JN. c-Myc is required for maintenance of glioma cancer stem cells. PLoS One. 2008;3:e3769. doi: 10.1371/journal.pone.0003769

65. Zheng H, Ying H, Yan H, Kimmelman AC, Hiller DJ, Chen AJ, Perry SR, Tonon G, Chu GC, Ding Z, et al. Pten and p53 converge on c-Myc to control differentiation, self-renewal, and transformation of normal and neoplastic stem cells in glioblastoma. Cold Spring Harb Symp Quant Biol. 2008;73:427–437. doi: 10.1101/sqb.2008.73.047

66. Schiapparelli P, Pirman NL, Mohler K, Miranda-Herrera PA, Zarco N, Kilic O, Miller C, Shah SR, Rogulina S, Hungerford W, et al. Phosphorylated WNK kinase networks in recoded bacteria recapitulate physiological function. Cell reports. 2021;36:109416. doi: 10.1016/j.celrep.2021.109416

67. Zhu W, Begum G, Pointer K, Clark PA, Yang SS, Lin SH, Kahle KT, Kuo JS, Sun D. WNK1-OSR1 kinase-mediated phospho-activation of Na+-K+-2Cl-cotransporter facilitates glioma migration. Mol Cancer. 2014;13:31. doi: 10.1186/1476-4598-13-31

68. Garzon-Muvdi T, Schiapparelli P, ap Rhys C, Guerrero-Cazares H, Smith C, Kim DH, Kone L, Farber H, Lee DY, An SS, et al. Regulation of brain tumor dispersal by NKCC1 through a novel role in focal adhesion regulation. PLoS Biol. 2012;10:e1001320. doi: 10.1371/journal.pbio.1001320

69. Haas BR, Cuddapah VA, Watkins S, Rohn KJ, Dy TE, Sontheimer H. With-No-Lysine Kinase 3 (WNK3) stimulates glioma invasion by regulating cell volume. Am J Physiol Cell Physiol. 2011;301:C1150–1160. doi: 10.1152/ajpcell.00203.2011

70. Moniz S, Martinho O, Pinto F, Sousa B, Loureiro C, Oliveira MJ, Moita LF, Honavar M, Pinheiro C, Pires M, et al. Loss of WNK2 expression by promoter gene methylation occurs in adult gliomas and triggers Rac1-mediated tumour cell invasiveness. Hum Mol Genet. 2013;22:84–95. doi: 10.1093/hmg/dds405

71. Hunter AW, Barker RJ, Zhu C, Gourdie RG. Zonula occludens-1 alters connexin43 gap junction size and organization by influencing channel accretion. Mol Biol Cell. 2005;16:5686–5698. doi: 10.1091/mbc.e05-08-0737

72. Rhett JM, Jourdan J, Gourdie RG. Connexin 43 connexon to gap junction transition is regulated by zonula occludens-1. Mol Biol Cell. 2011;22:1516–1528. doi: 10.1091/mbc.E10-06-0548

73. Grek CL, Montgomery J, Sharma M, Ravi A, Rajkumar JS, Moyer KE, Gourdie RG, Ghatnekar GS. A Multicenter Randomized Controlled Trial Evaluating a Cx43-Mimetic Peptide in Cutaneous Scarring. J Invest Dermatol. 2017;137:620–630. doi: 10.1016/j.jid.2016.11.006

74. Yamada K, Park HM, Rigel DF, DiPetrillo K, Whalen EJ, Anisowicz A, Beil M, Berstler J, Brocklehurst CE, Burdick DA, et al. Small-molecule WNK inhibition regulates cardiovascular and renal function. Nat Chem Biol. 2016;12:896–898. doi: 10.1038/nchembio.2168

75. Llombart V, Mansour MR. Therapeutic targeting of “undruggable” MYC. EBioMedicine. 2022;75:103756. doi: 10.1016/j.ebiom.2021.103756

76. Sun T, Warrington NM, Luo J, Brooks MD, Dahiya S, Snyder SC, Sengupta R, Rubin JB. Sexually dimorphic RB inactivation underlies mesenchymal glioblastoma prevalence in males. J Clin Invest. 2014;124:4123–4133. doi: 10.1172/JCI71048

77. Bayik D, Zhou Y, Park C, Hong C, Vail D, Silver DJ, Lauko A, Roversi G, Watson DC, Lo A, et al. Myeloid-Derived Suppressor Cell Subsets Drive Glioblastoma Growth in a Sex-Specific Manner. Cancer Discov. 2020;10:1210–1225. doi: 10.1158/2159-8290.CD-19-1355

78. Lee J, Nicosia M, Hong ES, Silver DJ, Li C, Bayik D, Watson DC, Lauko A, Kay KE, Wang SZ, et al. Sex-Biased T-cell Exhaustion Drives Differential Immune Responses in Glioblastoma. Cancer Discov. 2023;13:2090–2105. doi: 10.1158/2159-8290.CD-22-0869

