## Supplemental Figure Legends for "Connexin 43 drives glioblastoma cancer stem cell phenotypes through a WNK lysine-deficient protein kinase 1-c-MYC signaling axis"

**Supplemental Figure S1. Cx43 is required for GBM PDX CSC survival in the T3832 model.** (**A**) Schematic showing the nucleotide positions for each shRNA against *GJA1* (Cx43). (**B**) GBM PDX CSC line T3832 was subjected to lentiviral transduction with three non-overlapping shRNA constructs against *GJA1* (Cx43). The degree of knockdown was verified via immunoblotting using an antibody against Cx43. β-actin was used as a loading control. (**C**) Cell viability of T3832 CSCs containing shRNAs against Cx43 was measured via CellTiter-Glo on day 5 after plating. Luminescence values were normalized to day 0 values, and the fold change was calculated relative to the non-targeting control. n = 3 independent experiments, each performed in technical triplicate. (**D**) Apoptosis of T3832 CSCs containing shRNAs against Cx43 was measured via CaspaseGlo 3/7 on day 3 after plating. Luminescence values were normalized to cell number measured via CellTiter Glo at the same time point and then to the non-targeting control. n = 3 independent experiments, each performed in technical triplicate. (**E**) T3832 CSCs containing Cx43 shRNAs were plated in decreasing cell number (20, 10, 5, and 1 cells/well) with 24 replicates per number and evaluated for sphere formation 10-14 days later. n = 3 independent experiments. The online algorithm outlined in the Methods section was used to calculate stem cell frequency. *p < 0.05, **p < 0.01, ***p < 0.001 by one-way ANOVA analysis with Dunnett's Multiple Comparison Test.

**Supplemental Figure S2. Enrichment of MYC and MYC-related datasets in GBM cells expressing Cx43 shRNA.** GSEA was performed on differentially expressed genes that overlapped between DI318 expressing Cx43 sh73 and sh77. We focused on the TFT:TFT_LEGACY gene sets to gain mechanistic transcriptional information. Data sets related to MYC are boxed in red. The top 20 data sets are shown.

**Supplemental Figure 3. WNK1 is essential for the GBM PDX model T3832.** PDX CSC line T3832 was subjected to three different non-overlapping WNK1 shRNA transfections. (**A**) Knockdown was verified via immunoblotting using total WNK1 antibody. β-actin was used as a loading control. (**B**) Cell viability in T3832 containing *WNK1* shRNA was measured via CellTiter-Glo on day 5 after plating. n = 3 independent experiments, each performed in technical triplicate. (**C**) Cell death was measured in T3832 expressing *WNK1* shRNA via the CaspaseGlo 3/7 assay. n = 3 independent experiments, each performed in technical triplicate. (**D**) T3832 expressing *WNK1* shRNA were plated in decreasing cell number (20, 10, 5, 1 cell per well) with 24 replicates per condition and counted for sphere formation 10-14 days later. n = 3 independent experiments. The online algorithm outlined in the Methods section was used to calculate stem cell frequency. *$p < 0.05$, **$p < 0.01$, ***$p < 0.001$ by one-way ANOVA analysis with Dunnett's Multiple Comparison Test.

**Supplemental Figure S4. MYC is reduced after WNK1 shRNA.** (**A**) GSEA was performed on differentially expressed genes that overlapped between DI318 expressing WNK1 sh491 and sh920. We focused on the TFT:TFT_LEGACY gene sets to gain mechanistic transcriptional information. Data sets related to MYC are boxed in red. The top 20 data sets are shown. (**B**) MYC is also reduced in the T3832 model after depletion of WNK1 by shRNA. Actin was used as a loading control. n = 3 independent experiments. (**C**) mRNA expression of all WNK family members in DI318 as determined by RNA sequencing. n = 1 experiment performed in technical triplicate.
