## Supplementary figures and images for "Connexin 43 drives glioblastoma cancer stem cell phenotypes through a WNK lysine-deficient protein kinase 1-c-MYC signaling axis"

### Supplemental Figures

Supplemental Figure S1.

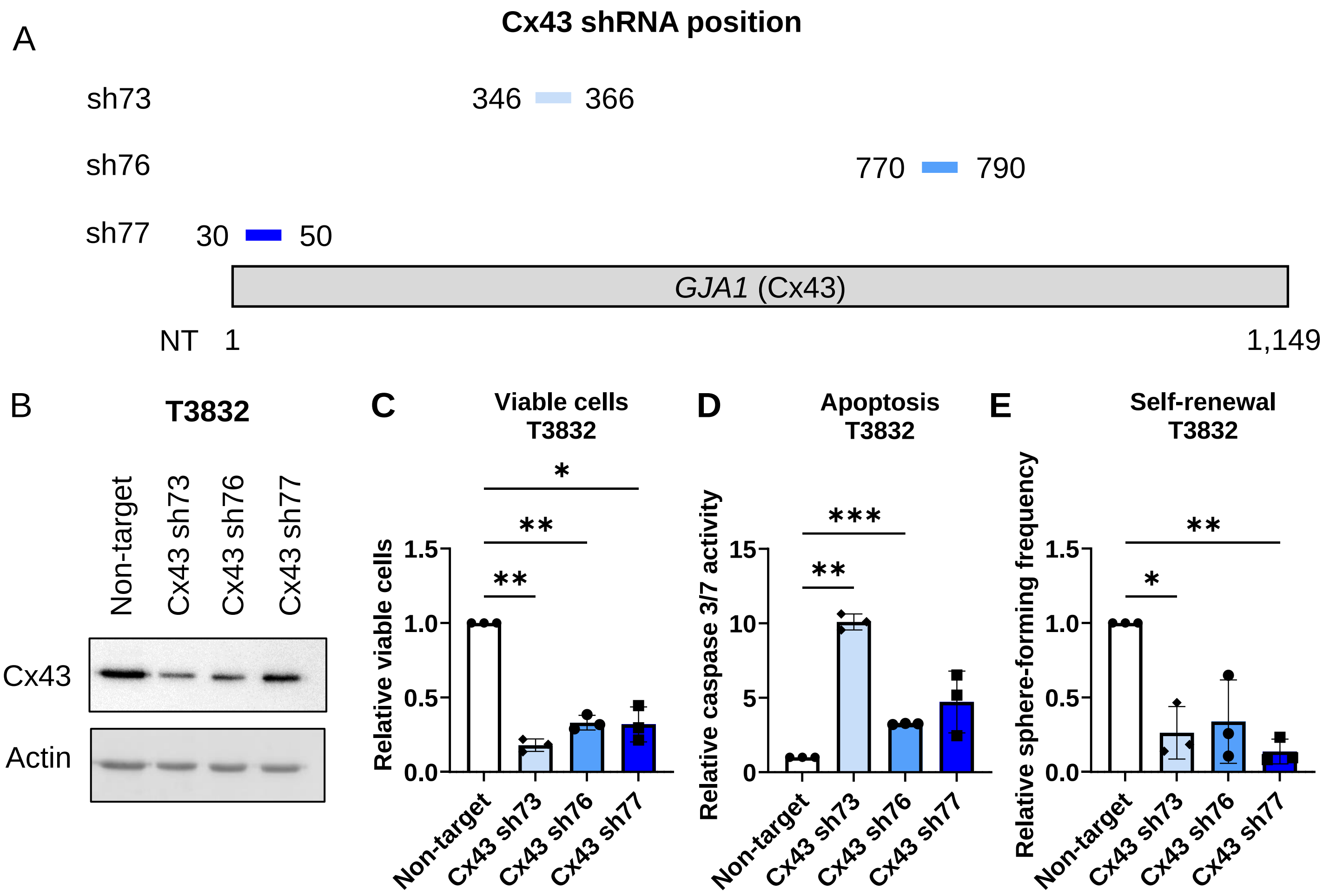

Supplemental Figure S2.

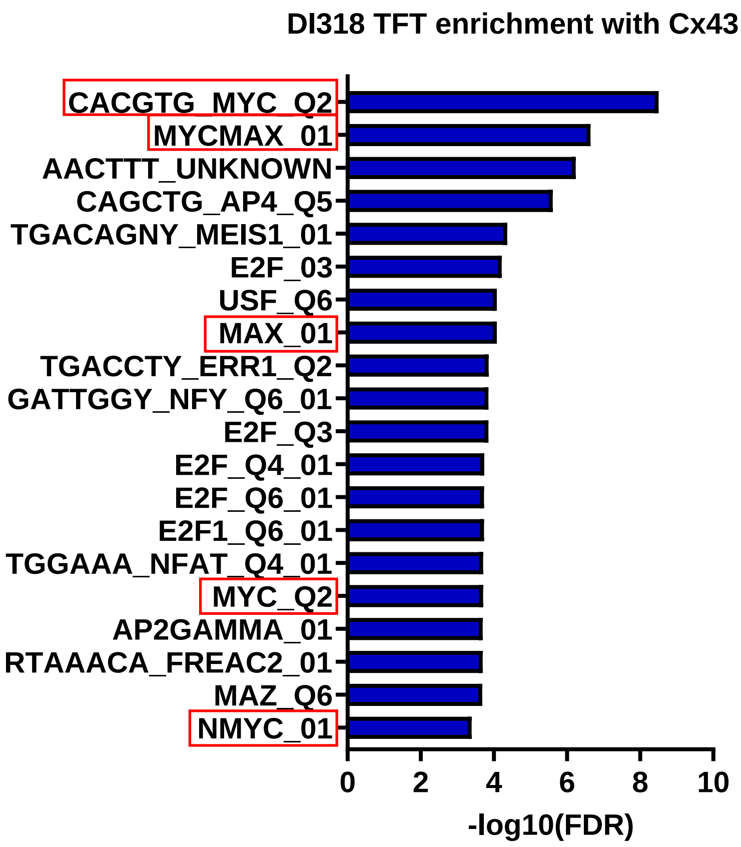

Supplemental Figure S3.

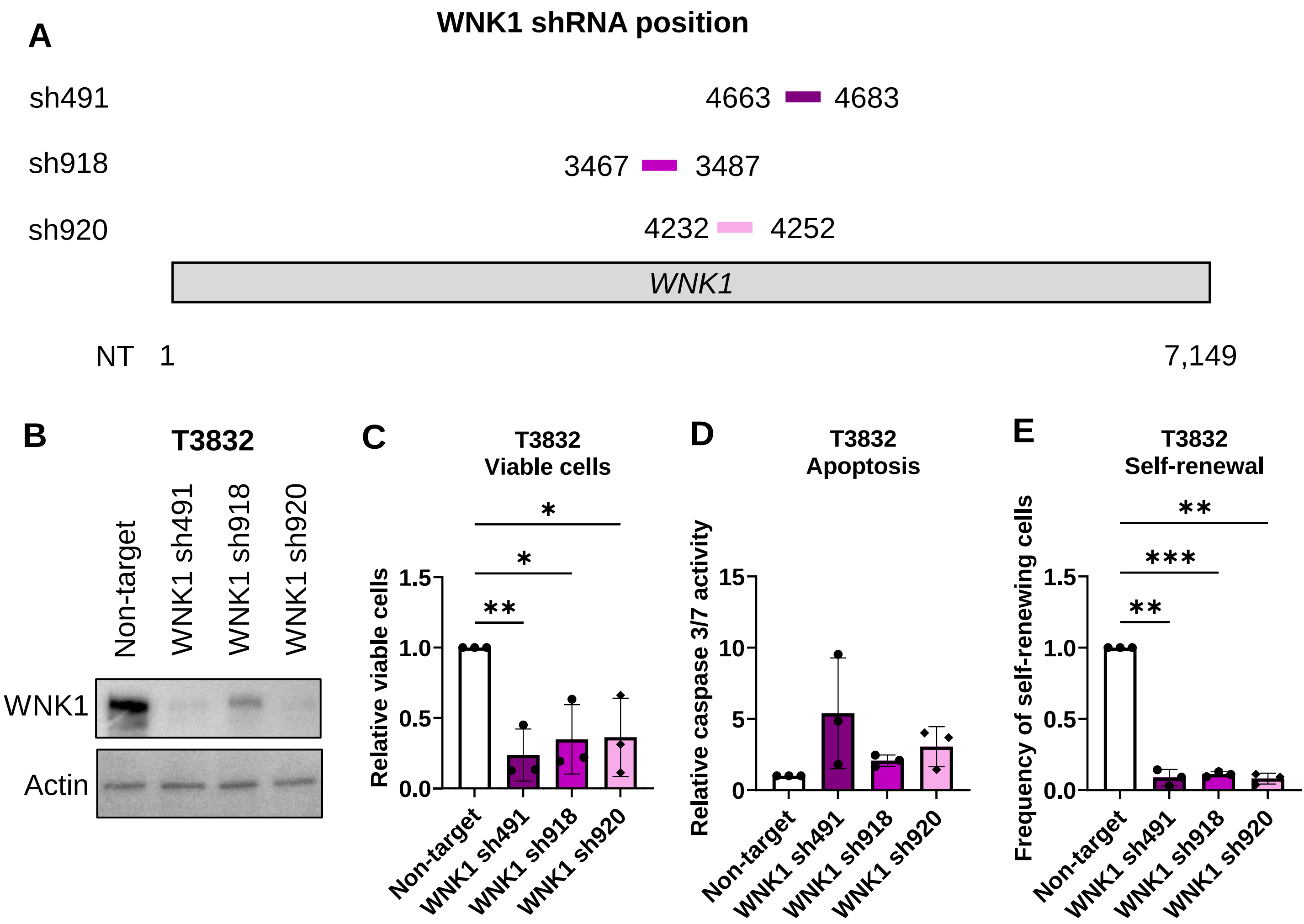

Supplemental Figure S4.

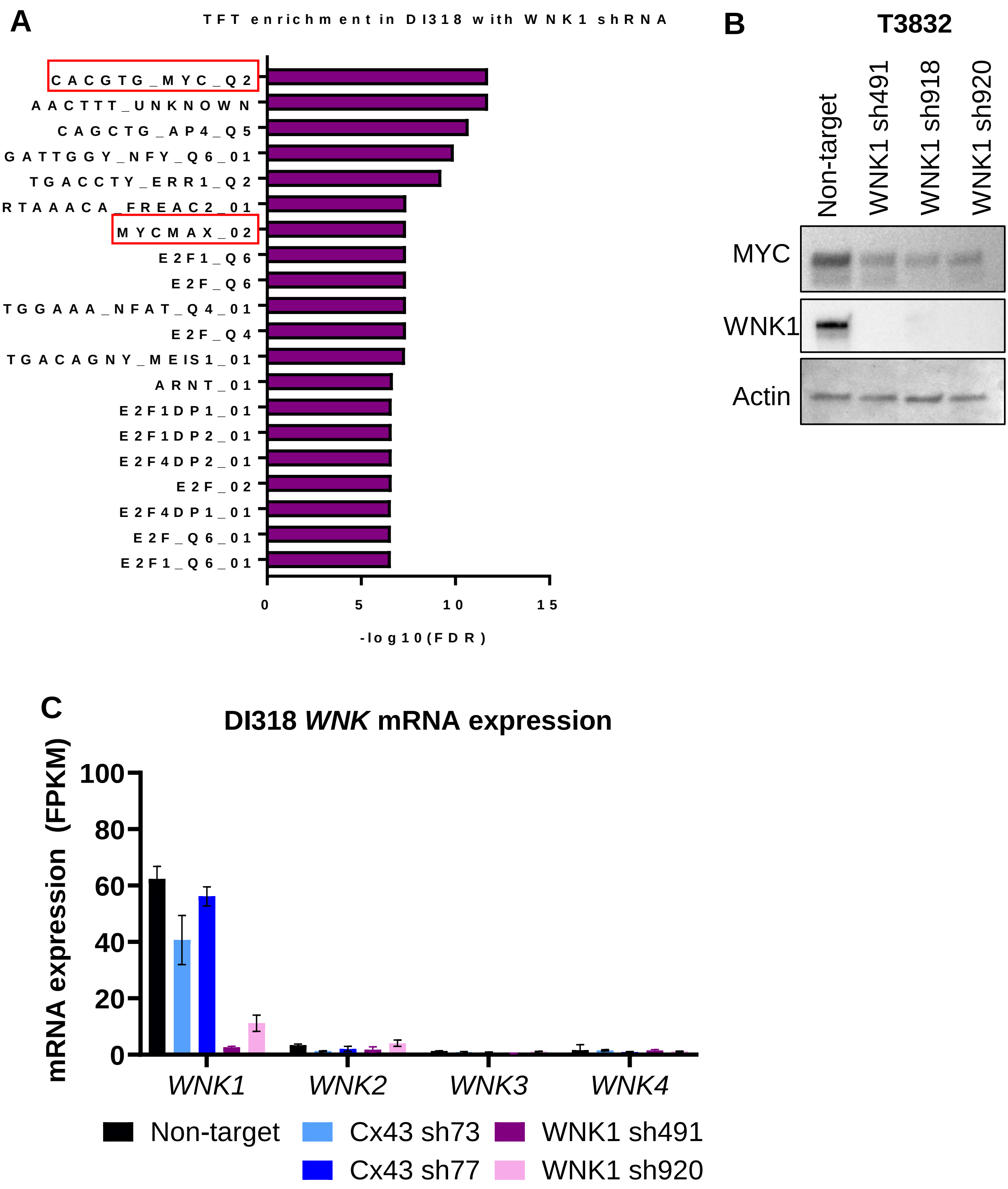
